# Glypican-3 (GPC3) is associated with MCPyV-negative status and impaired outcome in Merkel Cell Carcinoma

**DOI:** 10.1101/2022.02.06.479301

**Authors:** Sujatha Muralidharan, Thibault Kervarrec, Glen J. Weiss, Mahtab Samimi

## Abstract

**Background:** Merkel cell carcinoma (MCC) is an aggressive skin cancer, related to the Merkel Cell Polyomavirus (MCPyV) in 80% of cases. Immune checkpoint inhibitors provide sustained benefit in about half of MCC patients with advanced disease. Glypican-3 (GPC3) is an oncofetal tumor antigen that is an attractive target for chimeric antigen receptor T cell therapy due to its highly restricted expression on normal tissue and high prevalence in several solid tumors. GPC3 was previously found to be expressed in MCC but its association with tumor characteristics or prognosis has not been reported.

**Objectives:** To investigate the expression of GPC3 in MCC by immunohistochemistry (IHC) and its association with tumor characteristics, MCPyV status, and patient outcome.

**Methods:** The GC33 antibody clone was validated for GPC3 IHC staining of tumor specimens in comparison to an established GPC3 IHC antibody. A tissue microarray of tumors collected from an ongoing cohort of MCC patients was stained for GPC3 by IHC using GC33 antibody. Association of GPC3 positivity with baseline characteristics, MCPyV status (quantitative PCR) and outcome (death from MCC, recurrence) were assessed by Fisher’s exact tests and Cox regression analysis.

**Results:** Among 62 tumors from 59 patients, 42 samples (67.7%) were GPC3-positive. GPC3 expression was more frequently observed in females (p=0.048) and MCPyV-negative tumors (p=0.021). In the multivariate analysis, GPC3 expression was associated with increased death from disease (CSS) (hazard ratio [HR] 4.05, 95% CI 1.06-15.43), together with advanced age (HR 4.85, 95% CI 1.39-16.9) and male gender (HR 4.64, 95% CI 1.31-16.41).

**Conclusions:** GPC3 expression is frequently expressed in MCC tumors, especially MCPyV-negative cases, and is associated with increased risk of death. The high prevalence of surface GPC3 makes it a putative drug target.

## INTRODUCTION

Merkel cell carcinoma (MCC) is a rare and aggressive neuroendocrine skin cancer mostly occurring in elderly patients, with increased incidence in case of immunosuppression(1). MCC has a high propensity to metastasize, and the five-year overall survival has been estimated at 40%(2). While 80% of MCC are related to the integration of the oncogenic Merkel Cell Polyomavirus (MCPyV) genome into host cells(3), the remaining 20% cases are MCPyV-negative and harbor UV-induced mutations disrupting several oncogenic pathways(4). Both subsets were found to be immunogenic leading to the rationale of treating advanced stage MCC with immune checkpoint inhibitors (ICIs) such as PD-1/PD-L1 inhibitors. ICIs are considered a standard of care in patients with advanced disease(5,6). More than half of patients experience primary or secondary resistance to ICIs, which underlines the need for additional therapies, either as monotherapy or exerting synergistic effect with ICIs(7).

Glypican-3 (GPC3) is an oncofetal tumor antigen that is an attractive target for chimeric antigen receptor (CAR) T cell therapy due to its highly restricted expression on normal tissue and high prevalence in several adult and pediatric solid tumors(8). Aberrant GPC3 expression is implicated in tumorigenesis, and GPC3+ cancers are characterized by a highly immunosuppressive landscape which induces exhaustion in tumor-resident T cells(8). Drugs targeting GPC3 have been assessed in advanced cancer(9). GPC3 was previously found to be expressed in neuroendocrine small cell carcinomas including MCC(10), but its association with tumor stage, MCPyV status, or prognosis has not been characterized. The aim of the present study was to complete validation of a GPC3 antibody for use in immunohistochemistry (IHC), investigate the expression of GPC3 in MCC by IHC and to assess its association with tumor characteristics, MCPyV status, and patient outcome.

## MATERIAL AND METHODS

### Patients and tumor samples

MCC cases were selected from an ongoing historical/prospective cohort of MCC patients from France (local ethics committee approval, Tours, France, no. RCB2009-A01056-51) whose settings and inclusion criteria have previously been reported(11). Age, sex, American Joint Committee on Cancer (AJCC) stage at the time of diagnosis, immunosuppression (HIV infection, organ transplant recipients, hematological malignancies) and follow-up data (recurrence, death of any cause, death from MCC) were collected from patient files. Death was categorized as being related to MCC (CSS, cancer-specific death) or not (other cause) based on patients’ medical files. CSS was defined as the time from the initial confirmed diagnosis of MCC to the date of death related to MCC; overall survival (OS) as the time from diagnosis to the date of death regardless of cause; recurrence-free survival (RFS) as the time from diagnosis to the date of a clinical or paraclinical event related to MCC recurrence. Tumor samples had been collected during routine biopsies or surgeries as part of patients’ treatment plan.

Tumor samples were included in a tissue microarray (TMA), as previously described(11). Briefly, intratumor areas without necrosis were selected on hematoxylin phloxin saffron-stained before being extracted using a 1-mm tissue core, and cores were mounted in triplicate on the tissue microarray by using a semi-motorized tissue array system (MTA booster OI v2.00, Alphelys).

Twenty unique MCC whole mount specimen FFPE blocks (without clinical annotation) were sourced independently from AMS Bio. For GPC3 IHC validation with GC33 antibody, FFPE blocks for hepatocellular carcinoma (HCC), liposarcoma and non-small cell lung cancer (NSCLC) were sourced by NeoGenomics Laboratories, Inc. (Aliso Viejo, CA) from DLS and ProteoGenex. A normal tissue TMA consisting of different organ specimens from 3 unique individuals was sourced from US Biomax (Cat# FDA999w1).

### Determination of MCPyV status

MCPyV status was determined in MCC tumors using real time quantitative PCR as previously described. Briefly, genomic DNA was isolated from tumor samples and LTAg real-time PCR assay was performed using previously reported primers(11). Normalization was with albumin as the reference gene and the Waga MCC cell line (RRID:CVCL E998) was included as a control. The ΔCt method was used for quantification and results expressed as number of MCPyV copies/cells. MCPyV-positivity was defined for cases harboring MCPyV load >1.2 copies/cell(11).

### IHC assessment of GPC3

FFPE sections at 5μm were stained using anti-GPC3 mouse monoclonal primary antibody clone GC33 (Ventana Medical Systems, Inc. Tucson, AZ Cat# 790-45654) on BenchMark ULTRA to detect membrane and cytoplasmic expression. After anti-GPC3 primary antibody staining, heat-induced epitope retrieval was used followed by incubation of the primary antibody for 32 minutes. Immunodetection was accomplished with OptiView DAB Detection Kit (Ventana, Cat# 760-700). Isotype negative control and H&E staining was included for each specimen. Batch positive and negative controls were included on each stain run. Stained images were examined by a pathologist (MS) who was blinded to clinical data. In some cases, strong cytoplasmic staining complicated the ability to report strictly membrane staining as an individual compartment, therefore the final scoring of the tumor cells combined both cytoplasmic and membranous staining. Percent of tumor cells with GPC3 membrane and cytoplasmic staining at each intensity (0, 1+, 2+, 3+ corresponding to no staining, weak staining, moderate staining, and strong staining; respectively) were recorded and an overall H-score was calculated (range 0-300).

For validation of antibody clone GC33 for GPC3 IHC staining as a lab-developed test, all tumor tissue samples were stained with GC33 or 1G12 (a GPC3 IHC antibody clone used as an in vitro diagnostic and previously validated by NeoGenomics) and accuracy, sensitivity, specificity and precision of GC33 was determined compared to 1G12. Using comparisons between the 2 assays as a standard curve, a cutoff of H-score >30 for the GC33 was most comparable to a cutoff of H-score >20 as established for the 1G12 assay to report Positive or Negative for GPC3 expression (Figure S1). Specimens that showed consistent positive or negative results by both GC33 and 1G12 were considered true positive (TP) and true negative (TN) respectively. Any specimen reported positive by GC33 stain but negative by 1G12 or negative by GC33 but positive by 1G12 were considered false positive (FP) or false negative (FN) respectively. Accuracy was determined by comparing observed GC33 true staining to total specimens stained (TP + TN/Total specimens). Sensitivity was evaluated by comparing the observed GC33 true positive staining of the specimens to the expected positive expression (TP/TP+FN). Specificity was evaluated by comparing the observed GC33 true negative staining of the specimens to the expected negative expression (TN/TN+FP). Precision was determined by repeat staining of select specimens and examining concordance of results between repeat runs.

For MCC prevalence determination in terms of GPC3 protein expression by IHC, results from the validated GC33 assay were used. Interpretation of all immunostainings was blinded from clinicopathological parameters and patient outcome. When patients had more than one tumor sample included in the MCC TMA (for instance, primary tumor and metastasis), the GPC3 scoring of the primary tumor was included in the baseline and outcome analysis.

### Statistical analyses

Continuous data are described by medians (Q1–Q3) and categorical data with number and percentage of interpretable cases. Associations were assessed by two-tailed Fisher’s exact tests for categorical data. RFS, OS and CSS were analyzed by log-rank tests and presented as Kaplan-Meier curves. Univariate and multivariate Cox proportional-hazards regression was used to identify factors associated with outcome, estimating hazard ratios (HRs) and 95% confidence intervals (CIs). Covariates were identified as potential prognostic confounders with p≤0.25 on Cox univariate regression analysis and then included in the multivariate Cox analysis. P<0.05 was considered statistically significant. Statistical analysis involved use of XL-Stat-Life (Addinsoft, Paris, France).

## RESULTS

### Validation of GPC3 IHC using GC33 in MCC specimens

In order to validate the GPC3 IHC assay using GC33 antibody for use as a lab-developed test for clinical trials, TMA slides or whole mount slides containing tumor samples from HCC, liposarcoma, lung cancer, and MCC patients were stained by GPC3 IHC assays using either GC33 or previously validated 1G12 antibody clones. Scoring of these specimens based on GC33 vs. 1G12 staining was used to determine true and expected GPC3 expression (Table S1). These values were used to calculate accuracy, sensitivity, specificity and precision of GC33 staining for GPC3 compared to comparator 1G12 staining in tumor specimens for the purposes of GC33 assay validation. The GC33 assay was fully validated for use as a lab-developed test as it met all acceptable criteria (accuracy: 95%; sensitivity: 100%; specificity: 92%; precision: 100%) for each of these parameters and we used this assay for further assessments of MCC patient specimens.

### GPC3 expression in MCC and baseline characteristics

We examined MCC samples as well as normal tissue including skin for basal expression, using GPC3 GC33 IHC (**Fig. 1**). GPC3 was absent in normal skin. Among the 68 MCC tumors included in the TMA, 62 had at least one interpretable core for GPC3 staining (one core n=8, two cores n=25, three cores n=29). The median immunohistochemical GPC3 score was 85.8 (Q1-Q3 2.5-138.3, ranges 0-285) and accordingly, 42/62 samples (67.7%) were defined as GPC3-positive with an H-score >30. Representative IHC results are shown in **Fig. 1**. Among the 3 patients who had two tumor samples included in the TMA (primary tumor and lymph node metastasis), GPC3 expression was concordant between the primary tumor and the lymph node metastasis in two cases (both positive, n=1; both negative, n=1) and discordant in one case, where GPC3 expression was negative in the primary tumor and positive in the lymph node metastasis. Overall, 59 unique patients with MCC were included in the analysis reported in **Table 1**. GPC3 expression was more frequently expressed in female (27/35, 77%) than male patients (12/23, 52%) (p=0.048) and in MCPyV-negative (11/12, 92%) than MCPyV-positive tumors (26/44, 59%) (p=0.021). GPC3 expression was not associated with age, location of primary tumor, AJCC stage on diagnosis, type of tumor specimen (primary or metastasis), or immunosuppression (**Table 1**).

**Figure 1.**
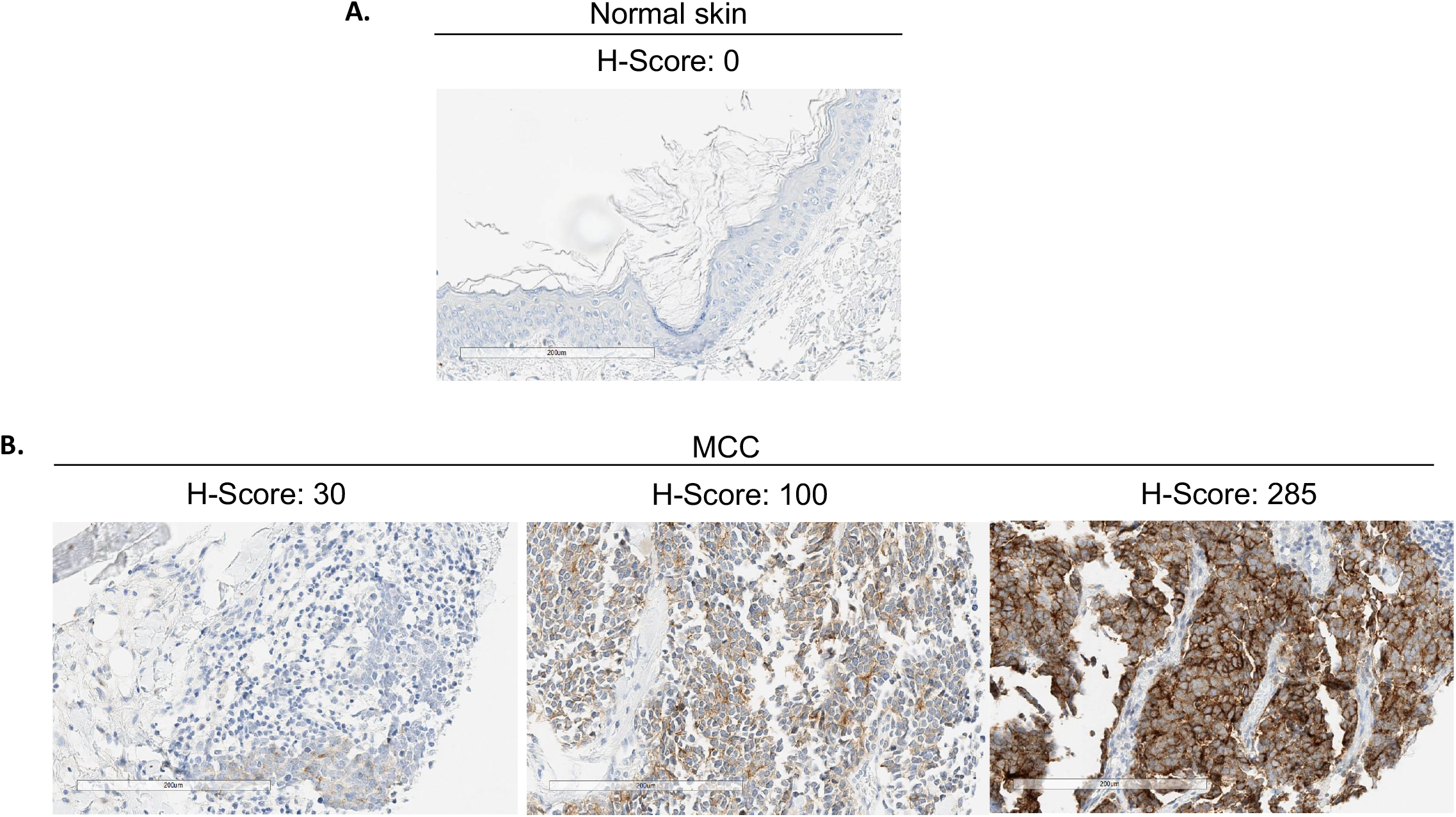
Representative GPC3 expression by IHC: Immunohistochemical staining of (**a**) normal skin and (**b**) MCC tumors for GPC3 expression. Representative images (20x magnification) with different levels of GPC3 (and H-scores) are shown. The scale bar represents 200 μm.

**Table 1.** GPC3 expression according to baseline characteristics.

| Characteristics | All patients | GPC3 expression |  | P-value<br>(Fisher's exact<br>test) |
| --- | --- | --- | --- | --- |
|  | (n=59)<br>(n,%) | Negative (n=20)<br>(n,%) | Positive (n=39)<br>(n,%) |  |
| **Age (\*)** |  |  |  | 0.780 |
| **> median (80.5 y)** | 29 (50.0) | 10 (52.6) | 19 (48.7) |  |
| **< median (80.5 y)** | 29 (50.0) | 9 (47.4) | 20 (51.3) |  |
| **Sex (\*)** |  |  |  | **0.048** |
| **Female** | 35 (60.3) | 8 (42.1) | 27 (69.2) |  |
| **Male** | 23 (39.7) | 11 (57.9) | 12 (30.8) |  |
| **AJCC stage (at diagnosis)** |  |  |  | 0.162 |
| **(\*)** | 20 (34.5) | 8 (42.1) | 12 (30.8) |  |
| **Stage 1** | 19 (32.8) | 3 (15.8) | 16 (41.0) |  |
| **Stage 2** | 18 (31.0) | 8 (42.1) | 10 (25.6) |  |
| **Stage 3** | 1 (1.7) | 0 (0.0) | 1 (2.6) |  |
| **Stage 4** |  |  |  |  |
| **Immunosuppression (\*)** |  |  |  | 0.975 |
| **No** | 52 (89.7) | 17 (89.5) | 35 (89.7) |  |
| **Yes** | 6 (10.3) | 2 (10.5) | 4 (10.3) |  |
| **Location of primary (\*)** |  |  |  | 0.897 |
| **head and neck** | 19 (32.8) | 5 (26.3) | 12 (30.8) |  |
| **limb** | 28 (48.3) | 10 (52.6) | 16 (41.0) |  |
| **trunk** | 5 (8.6) | 2 (10.5) | 10 (25.6) |  |
| **occult primary** | 6 (10.3) | 2 (10.5) | 1 (2.6) |  |
| **Type of sample** |  |  |  | **0.594** |
| **Primary** | 41 (69.5) | 13 (65.0) | 28 (71.8) |  |
| **Metastasis** | 18 (30.5) | 7 (35.0) | 11 (28.2) |  |
| **MCPyV status (\*\*)** |  |  |  | **0.021** |
| **Negative** | 12 (21.4) | 1 (5.3) | 11 (29.7) |  |
| **Positive** | 44 (78.6) | 18 (94.7) | 26 (70.3) |  |

*(\*) data missing for one patient ; (\*\*) data missing for 3 patients*

**Table 2.** Univariate and multivariate Cox proportional hazard analysis for MCC related death, death from any cause et MCC recurrence.

| Covariate | Cancer Specific survival |  |  |  | Overall survival |  |  |  | MCC Recurrence (any site) |  |  |  |
| --- | --- | --- | --- | --- | --- | --- | --- | --- | --- | --- | --- | --- |
|  | Univariate analysis |  | Multivariate analysis |  | Univariate analysis |  | Multivariate analysis |  | Univariate analysis |  | Multivariate analysis |  |
|  | HR (95% CI) | p | HR (95% CI) | p | HR (95% CI) | p | aHR (95% CI) | p | HR (95% CI) | p | aHR (95% CI) | p |
| **Age (years)** |  |  |  |  |  |  |  |  |  |  |  |  |
| >80.5 vs ≤ 80.5 | 2.86 (1.08-7.54) | 0.034 | **4.85 (1.39-16.9)** | **0.013** | 2.85 (1.37-5.95) | 0.005 | **3.85 (1.56-9.48)** | **0.003** | 1.13 (0.50-2.56) | 0.772 | 0.75 (0.26-2.16) | 0.590 |
| **Sex** |  |  |  |  |  |  |  |  |  |  |  |  |
| male vs female | 1.99 (0.81-4.92) | 0.135 | **4.64 (1.31-16.41)** | **0.017** | 1.43 (0.71-2.88) | 0.321 | **2.75 (1.10-6.84)** | **0.030** | 1.94 (0.85-4.42) | 0.113 | 1.56 (0.57-4.28) | 0.389 |
| **AJCC** |  |  |  |  |  |  |  |  |  |  |  |  |
| 3-4 vs 1-2 | 1.82 (0.73-4.55) | 0.196 | 2.16 (0.73-6.35) | 0.161 | 0.90 (0.42-1.96) | 0.800 | 1.65 (0.68-3.98) | 0.265 | 1.41 (0.60-3.34) | 0.431 | 1.64 (0.63-4.28) | 0.314 |
| **Immunosuppression** |  |  |  |  |  |  |  |  |  |  |  |  |
| present vs absent | 1.85 (0.54-6.41) | 0.327 | 3.29 (0.79-13.71) | 0.101 | 3.23 (1.28-8.14) | 0.013 | **5.25 (1.82-15.2)** | **0.002** | 1.39 (0.41-4.70) | 0.593 | 2.24 (0.60 – 8.34) | 0.230 |
| **MCPyV status** |  |  |  |  |  |  |  |  |  |  |  |  |
| positive vs negative | 0.26 (0.00-0.67) | 0.005 | 0.42 (0.16-1.09) | 0.076 | 0.29 (0.00-0.61) | 0.001 | **0.37 (0.00-0.83)** | **0.017** | 0.23 (0.00-0.55) | 0.001 | **0.20 (0.00 – 0.57)** | **0.002** |
| **GPC3 expression** |  |  |  |  |  |  |  |  |  |  |  |  |
| positive vs negative | 2.24 (0.74-6.75) | 0.153 | **4.05 (1.06-15.43)** | **0.040** | 1.58 (0.72-3.46) | 0.252 | 2.42 (0.89-6.54) | 0.082 | 1.45 (0.60-3.53) | 0.412 | 1.34 (0.48-3.79) | 0.576 |

*HR, Hazard Ratio, aHR, adjusted HR ; CI, confidence interval ; MCC, Merkel Cell Carcinoma*

### Independent cohort of MCC samples

In evaluation of the independent cohort of whole mount MCC samples (n=20) with the GPC3 GC33 IHC assay, 14 samples showed positive GPC3 expression (H score >30) indicating a prevalence of 70% GPC3 positive cases in MCC. This was consistent with the GPC3+ prevalence determination based on MCC cases from the TMA.

### GPC3 expression and patient outcome

Data were available for 59 patients, with a median follow up of 87.7 months (95% CI 74.0-97.5). Mean OS was 67.7 months (95% CI 53.7-81.7) and mean CSS was 85.0 months (95% CI 70.3-99.6). During follow up, 23 patients had recurred (38.9%) and 32 patients (54.2%) had died including 19 from MCC (59.3%). As shown in **Fig. 2**, five-year CSS was numerically higher in GPC3-negative than GPC3-positive patients (77.2%, 95%CI 57.3-97.0 vs 57.1%, 95% CI 40.6-73.6) (log-rank test, p=0.142). OS and RFS did not differ significantly between groups (**Fig. 2**). In the multivariate analysis, GPC3 expression was associated with worse CSS (HR 4.05, 95% CI 1.06-15.43), together with advanced age (HR 4.85, 95% CI 1.39-16.9) and male gender (HR 4.64, 95% CI 1.31-16.41). By contrast, MCPyV-positive MCC cases were associated with reduced risk of death (HR 0.37, 95% CI 0.00-0.83) and reduced risk of recurrence (HR 0.20, 95% CI 0.00 – 0.57). GPC3 had no significant association with OS, while advanced age (HR 3.85, 95% CI 1.52-9.48), male gender (HR 2.75, 95% CI 1.10-6.84), and immunosuppression (HR 5.25, 95% CI 1.82-15.2) were associated with worse OS.

**Figure 2.**
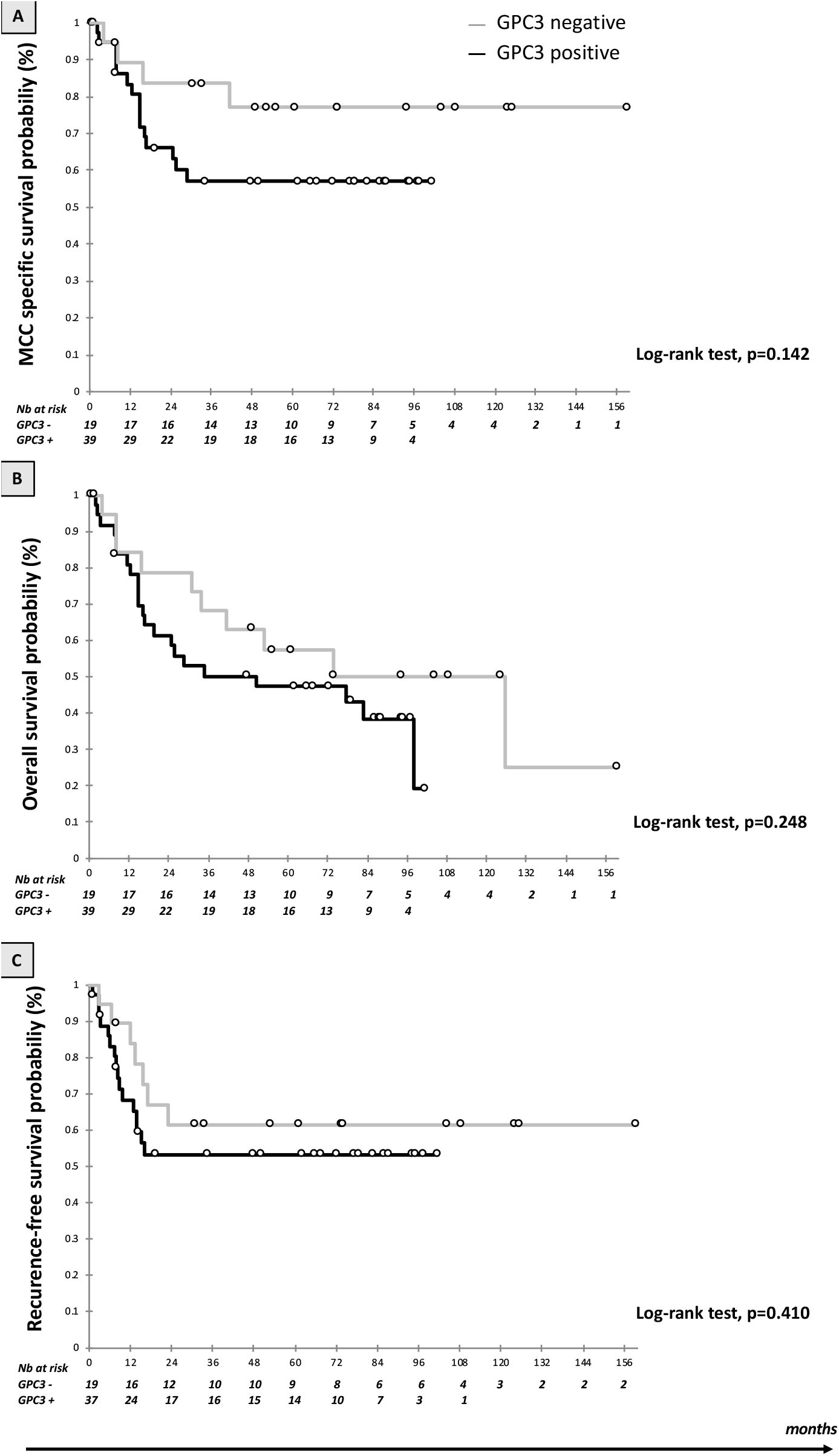
Kaplan Meier curves according to GPC3 expression: (**a**) Survival estimates of cancer-specific survival (CCS); (**b**) Survival estimate of overall survival (OS); and (**c**) Survival estimates of recurrence-free survival (RFS). Black bar: Positive GPC3 expression and grey bar: negative GPC3 expression.

## DISCUSSION

In this study, we found that GPC3 was expressed in nearly 70% of MCC tumors and up to 90% of MCPyV-negative cases, and was associated with worse prognosis in terms of risk of death from MCC. GPC3 is an heparan sulfate proteoglycan which is expressed in embryonic tissues and solid cancers such as hepatocellular carcinoma, ovarian clear cell carcinoma, melanoma, squamous cell carcinoma of the lung, and some pediatric cancers(9). GPC3 was not expressed in normal skin in our samples, similar to other results(12). Our results are in line with one previous study which reported increased GPC3 in MCC at the transcriptional level(13). Another study reported GPC3 expression by IHC in 39 out of 55 MCC cases (71%), without providing details on tumor characteristics or clinical outcomes(10). In our study, GPC3 expression was associated with death from MCC, in line with reports of its prognostic role in hepatocellular carcinoma(14). Indeed, GPC3 was reported to participate in tumor growth and promote epithelial-mesenchymal transition by impacting several signaling pathways, such as upregulation of the Wnt signaling, ERK pathway, YAP and hedgehog cascades(15). In the setting of MCC, the significant differential expression with MCPyV-positive and MCPyV-negative remains elusive, given that little is known on the regulation of GPC3 expression. Although no *GPC3* mutations have been identified so far in MCPyV-negative MCCs, GPC3 was shown to be a transcriptional target of c-Myc in the setting of hepatocellular carcinoma(16). Interestingly, *MYC* family gene amplification and MYC protein expression have previously been reported in virus-negative MCCs(17–19). On the other hand, the MCPyV oncoprotein ST forms a complex with the MYC paralog MYCL (L-MYC) and its heterodimeric partner MAX, the complex recruiting EP400 chromatin remodeling complex, which in turn bind to the transcriptional start sites of several hundred target genes, which encompass a large number of known MYC target genes(20).

Limitations to our study include a small cohort of predominantly early-stage MCC and that this was a retrospective analysis. Additional independent clinical prevalence studies in advanced MCC are needed to validate our findings.

In conclusion, GPC3 expression, advanced age, and male gender were associated with impaired CSS in MCC in this study, whereas MCPyV-positivity was significantly associated with reduced risk of death and recurrence. Given its high prevalence in MCC and restricted expression in normal tissue, GPC3 is an attractive target for CAR T cell therapy. The GC33 clone performs similarly to 1G12 and will be used for IHC screening for clinical trials. A new CAR T trial evaluating a GPC3 directed CAR T in patients with solid tumors including MCC is due to activate (NCT05120271).

## Acknowledgements

Authors acknowledge dermatologists and pathologists who included patients in the MCC cohort (Drs. Guido Bens, Eric Estève, and Patrick Michenet, CHR Orléans, France; Drs. Yannick Le Corre and Sophie Michalak-Provost, CHU Angers, France; Prof. François Aubin and Dr. Charlée Nardin, CHU Besançon, France; Drs. Ewa Wierzbiecka-Hainault and Eric Frouin, CHU Nantes, France; Prof. Philippe Saïag and Dr. Astrid Blom, Hospital Ambroise Paré, Paris, France; and Anne Tallet (Plateforme de Génétique Moléculaire des Cancers, CHU Tours, France) who performed the qPCR analyses.

## FIGURE LEGENDS

**Figure S1. Comparisons between the 2 IHC assays**. Legend: x-axis is GC33, y-axis is 1G12, best fit lit is plotted with an r^2^ co-efficient of 0.9329.

