## Supplementary figures and images for "Glypican-3 (GPC3) is associated with MCPyV-negative status and impaired outcome in Merkel Cell Carcinoma"

### Figure S1

#### Slide 1
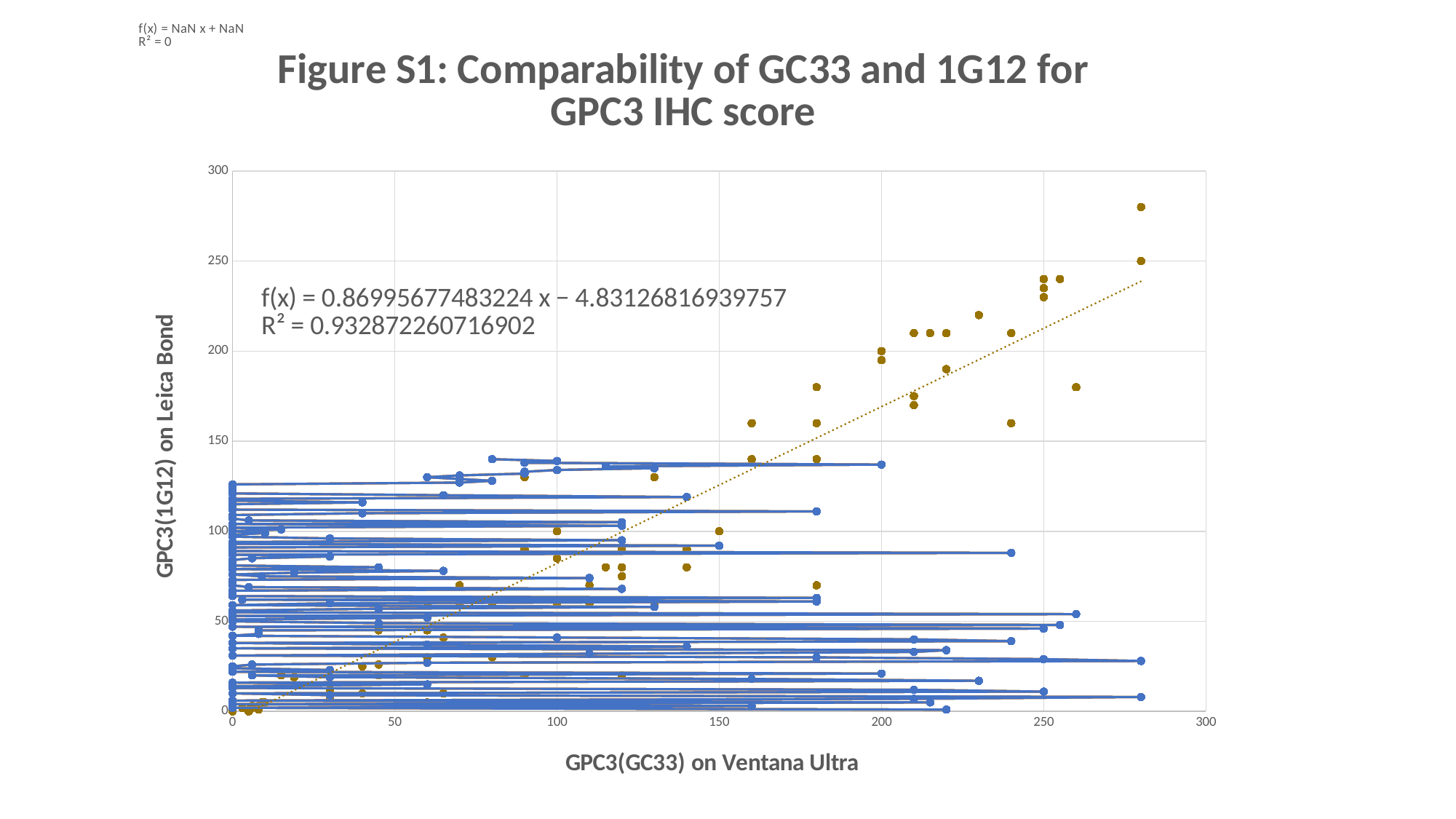

##### Chart: Figure S1: Comparability of GC33 and 1G12 for GPC3 IHC score
| Category |
|---|

### Table S1

#### Slide 1
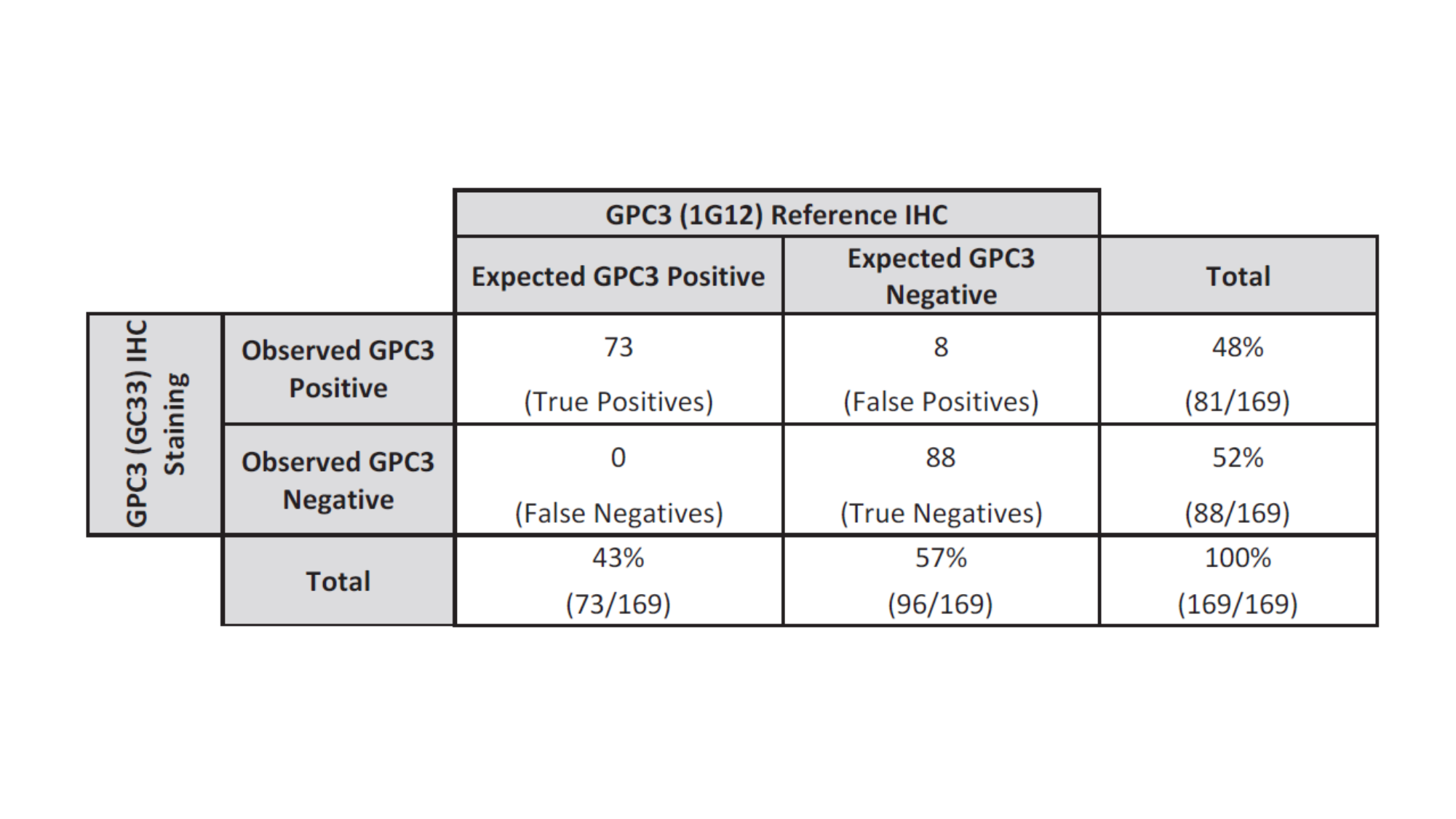
